# *Tbx1* interacts genetically with *Vegfr3* to regulate cardiac lymphangiogenesis in mice

**DOI:** 10.1101/553578

**Authors:** Stefania Martucciello, Maria Giuseppina Turturo, Sara Cioffi, Li Chen, Antonio Baldini, Elizabeth Illingworth

## Abstract

The transcription factor *TBX1* is the major gene implicated in 22q11.2 deletion syndrome. The complex clinical phenotype includes vascular anomalies and a recent report presented new cases of primary lymphedema in 22q11.2DS patients. We have previously shown that Tbx1 activates Vegfr3 gene expression in lymphatic endothelial cells and that this activation is critical for lymphatic vessel development in prenatal mice and for their survival post-natally. Using loss-of-function genetics and transgenesis, we show a strong genetic interaction between *Tbx1* and *Vegfr3* in cardiac lymphangiogenesis that causes cardiac lymphatic vessel anomalies in compound heterozygotes. Intriguingly, different aspects of the cardiac lymphatic phenotype were regulated independently by the two genes. *Tbx1*^*Cre*^-activated Vegfr3 transgene expression was able to rescue the morphological abnormalities in the cardiac lymphatic vessels of compound heterozygotes, but it did not rescue the severe cardiac lymphatic vessel hypoplasia observed in *Tbx1* homozygotes. Moreover, our study revealed a differential sensitivity between the ventral and dorsal cardiac lymphatic networks to the effects of altered *Tbx1* and *Vegfr3* gene dosage. Overall, our study demonstrates that a fine dosage balance between *Tbx1* and *Vegfr3* is required to regulate the number and morphology of cardiac lymphatic vessels.

## INTRODUCTION

*Tbx1* is a major developmental gene whose critical role in lymphangiogenesis has been demonstrated in the mouse (1). In mice, in the absence of *Tbx1*, most lymphatic vessels are lost around embryonic day (E)16.5. We have shown that Tbx1 regulates *Vegfr3* expression by binding to an intragenic enhancer element in the endogenous *Vegfr3* gene. In *Tbx1* germline mutants and in EC-specific *Tbx1* conditional mutants, *Vegfr3* expression in lymphatic ECs is lost between E15.5 and 16.5, which led us to hypothesize that lymphatic vessels in *Tbx1* mutants are not maintained because of reduced *Vegfr3* expression in lymphatic ECs (1).

In this study, we have investigated the interplay between Tbx1 and Vegfr3 in cardiac lymphangiogenesis by manipulating the dosage of both genes in ECs. We focused our attention on cardiac lymphangiogenesis because the cardiac lymphatic phenotype is severe in *Tbx1* mutants (1) and because cardiac neo-lymphangiogenesis has been implicated in the response to ischemia (2) (3) (4).

In the mouse, the cardiac lymphatic network develops from lymphatic ECs that migrate from extra-cardiac tissues into the heart around E12.5. As embryonic development progresses, an extensive network of sub-epicardial lymphatic vessels forms, extending over the heart from base to apex and covering mainly the left ventricle, both ventrally and dorsally, while fewer lymphatic vessels extend over the right ventricle (5)(2). The cardiac lymphatic vessel network is complete at postnatal day (P) 15.

As in other organs, in the heart, the primary role of the lymphatic vessels is to maintain fluid homeostasis and transport immune cells. In the adult heart, tissue fluid absorbed by the lymphatic capillaries drains into two main pre-collector vessels within the heart, that run along the left conal vein dorsally, and the left cardiac vein ventrally, and ultimately into the mediastinal lymph nodes. Studies in mice show that acute inflammation provokes neolymphangiogenesis, which is important for antigen clearance and for the resolution of inflammation (6)(7). Recently, Klotz et al., showed that cardiac injury (myocardial infarction) provokes a robust neolymphangiogenic response and upregulation of developmental lymphatic genes (*Vegfr3, Prox1, Lyve1*). Furthermore, post-infarct cardiac function improved after treatment with recombinant Vegf-C (2), suggesting that the Vegf-C-Vegfr3 axis is important for responding to pathological conditions of the heart.

In this study, we use genetic experiments to demonstrate a strong interaction between Tbx1 and Vegfr3 in cardiac lymphangiogenesis. Increasing endothelial expression of *Vegfr3* was sufficient to rescue partially the cardiac lymphatic phenotype in compound heterozygous mutants. However, it did not improve cardiac lymphatic development in *Tbx1* homozygous mutants, which lack lymphatic vessels in most tissues, including the heart. Overall, our data suggest that a dosage balance between Tbx1 and Vegfr3 is critical for cardiac lymphatic development.

## RESULTS

### *Tbx1* and*Vegfr3* expression overlap in cardiac lymphatics

We used a *lacZ* reporter allele (8) and monitored *Tbx1* expression by β-galactosidase (β-gal) activity in *Tbx1*^*lacz*/+^ hearts at E18.5 (Fig. 1). This revealed β-gal+ vessel-like structures on the surface of the heart that resembled lymphatic vessels (Fig. 1A, a, a’). To determine whether these were in fact lymphatic vessels, we immunostained the hearts of wild type littermates with anti-Vegfr3 antibody, which specifically labels lymphatic endothelial cells at this developmental stage (9). Results indicated that *Tbx1* (Fig. 1A, a, a’) and *Vegfr3* (Fig. 1A, b, b’) had near identical expression, indicating that they are co-expressed in most cardiac lymphatic vessels at this stage of development. Note that *Tbx1* is also expressed in the conotruncus (arrow in Fig. 1A, a), as previously reported (10) (11).

**Figure 1.**
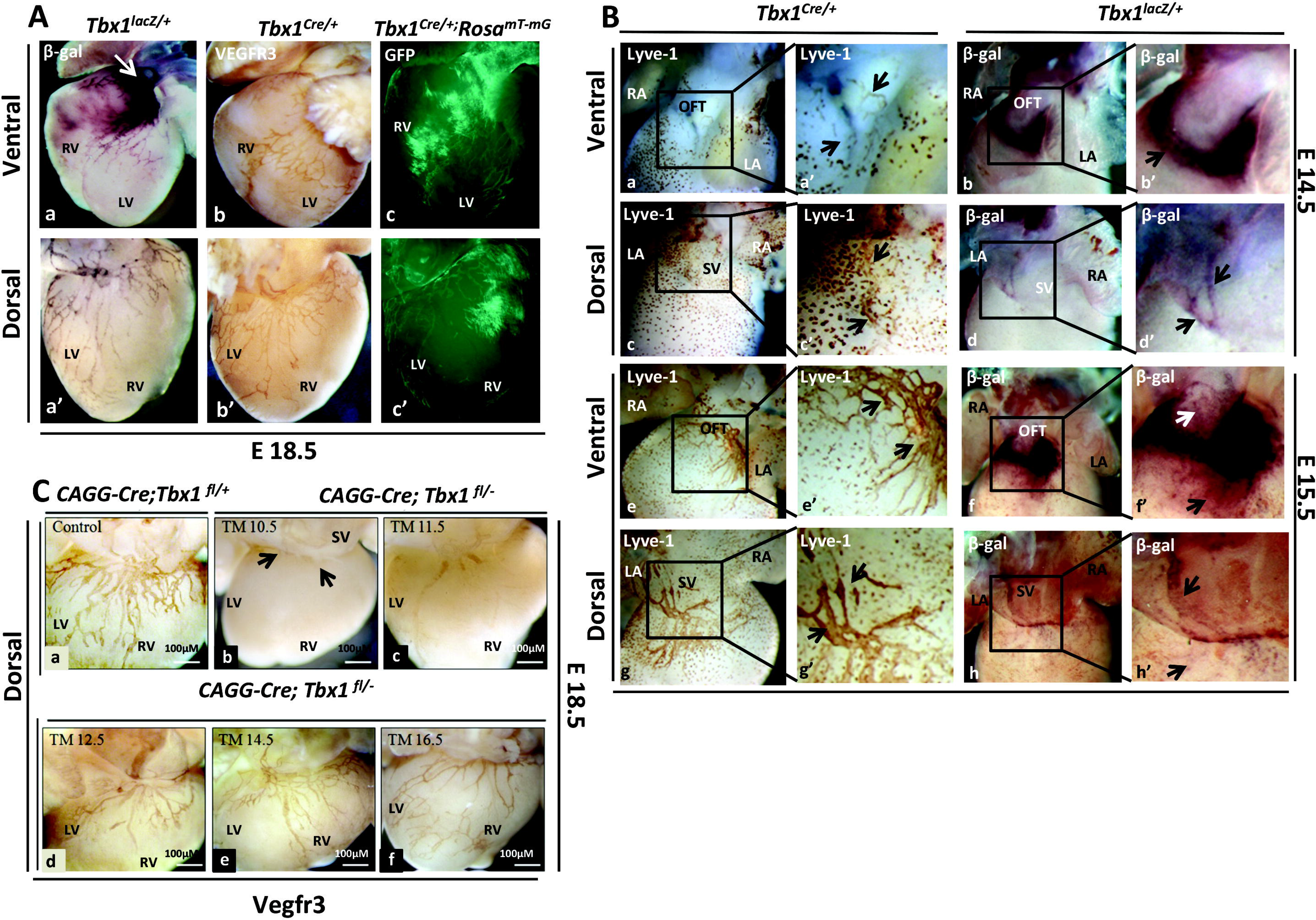
*Tbx1 and Vegfr3 are co-expressed in cardiac lymphatic vessels.* (A) β-gal staining of E18.5 *Tbx1*^*lacZ*/+^ hearts shows that *Tbx1* is expressed in the conotruncus (arrow in a) and in cardiac lymphatic vessels (a-a’), identified in b-b’ by anti-VEGFR3 immunostaining. Fate mapping of *Tbx1*-expressing cells and their descendents in *Tbx1*^*cre*/+;^ *Rosa*^*mT-mG*^ hearts (c-c’) reveals a contribution to cardiac lymphatic vessels and to the right ventricular myocardium. (B). Vegfr3 immunostaining (left) of E.14.5 (a-d) and E.15.5 (e-h) *Tbx1*^*Cre*/+^ hearts reveals early cardiac lymphatic vessel development (arrows in a’, c’, e’, g’). β-gal staining (right) of E.14.5 (b, b’, d, d’) and E.15.5 (f, f’, h, h’) *Tbx1*^*lacZ*/+^ hearts shows that *Tbx1* is expressed in lymphatic vessels at the base of the outflow tract ventrally (arrows in b’, f’) and in the sinus venosus dorsally (arrows in d’, h’). *Temporal requirement of Tbx1 for cardiac lymphatic vessel development* (C). Cardiac lymphatic vessels revealed by Vegfr3 immunostaining of E18.5 hearts following Tamoxifen (TM) induced inactivation of *Tbx1* between E10.5 and E16.5 (Fig. 1C, b-f) shows that Tbx1 is required for cardiac lymphangiogenesis before E14.5. Scale bar 100μM Abbreviations: LV, left ventricle, RV, right ventricle, SV, sinus venosus, OFT, outflow tract.

In order to understand where Tbx1 and Vegfr3 may interact to promote cardiac lymphangiogenesis, we first mapped the distribution of *Tbx1*-expressing cells and their descendants in embryonic mouse hearts, and then immunostained the same hearts with anti-Vegfr3. For this, we crossed *Tbx1*^*Cre*/+^ mice with R26R-*lacZ* reporter mice (12) and evaluated *Tbx1*-induced recombination in the hearts of *Tbx1*^*Cre/+;*^*R26R* embryos at E18.5 by β-gal staining. The hearts were then immunostained with anti-Vegfr3 antibody. Results revealed extensive overlap between *Vegfr3* expression and β-gal staining in cardiac lymphatic vessels (Supplementary Fig. S1), indicating that *Tbx1*-expressing cells and their descendants populate the majority of cardiac lymphatic vessels. Furthermore, the lack of β-gal+; Vegfr3-negative vessels indicates that in the mouse heart, they contribute exclusively to lymphatic vessels and not to veins and arteries. *Tbx1*^*Cre*^ activates recombination in cardiac muscle of the right ventricle but this tissue does not express *Vegfr3,* thereby excluding this tissue as a potential site of interaction. We also evaluated the distribution of *Tbx1*-expressing cells and their descendants using an alternative reporter (*Rosa*^*mT-mG*^) (13). For this, we crossed *Tbx1*^*Cre*/+^ mice with *Rosa*^*mT-mG*^ mice and analyzed the hearts of *Tbx1*^*Cre/+;*^ *Rosa*^*mT-mG*^ embryos at E18.5, where *Tbx1*-expressing cells and their descendants were labelled by *Tbx1*^*Cre*^-activated expression of a fluorescent (GFP) reporter (Fig. 1A, c, c’). Results were comparable to those obtained with the R26R-*lacZ* reporter.

In the mouse, the first cardiac lymphatic vessels form around E14.5 at the sinus venosus on the dorsal surface of the heart and at the base of the aorta on the ventral surface, from where they extend in a basal to apical direction (5)(2). We asked what is the earliest time point of *Tbx1* expression in cardiac lymphatic vessels? To address this question we performed β-gal staining and Lyve1 immunostaining on *Tbx1*^*lacz*/+^ hearts at E14.5 and E15.5 (anti-Lyve1 labels lymphatic ECs). Results showed that at E14.5 (Fig. 1B, a-d) and E15.5 (Fig. 1B, e-h), on the dorsal surface of the heart, lymphatic vessels (Lyve1+) at the sinus venosus (Fig. 1B, c, c’, g, g’) also expressed *Tbx1* (β-gal+) (arrows in d’ and h’). This was less evident on the ventral surface of the E14.5 heart, because *Tbx1* is highly expressed in the conotruncus (arrow in b’), where the first ventral lymphatic vessels form (arrows in a’). However, at E15.5, a small network of *Tbx1*-expressing lymphatic vessels was visible on the ventral surface of the outflow tract (white arrow in f’) and heart (black arrow in f’). Thus, we conclude that *Tbx1* is expressed in the earliest forming cardiac lymphatic vessels.

### *Tbx1* is expressed and required in the early phases of cardiac lymphatic development

In order to pinpoint the critical time requirement for *Tbx1* in cardiac lymphatic development, we crossed TgCAGG-CreER™;*Tbx1*^*lacZ*/+^ mice with *Tbx1*^*flox/flox*^ mice and inactivated *Tbx1* in a time-specific manner, as previously described (10). We injected pregnant female mice with a single dose of tamoxifen (TM) at E10.5, E11.5, E12.5, E14.5 and E16.5. We then harvested embryos at E18.5 and analyzed the cardiac lymphatic vessels by Vegfr3 immunostaining (Fig. 1C). Results showed that after injection of TM at E10.5 (Fig. 1C, b) lymphatic vessels were present on the surface of the sinus venosus (arrows in b) but not in the ventricular myocardium. TM injection at E11.5 and E12.5 (Fig. 1C, c, d) resulted in cardiac lymphatic vessel hypoplasia, while after TM injection at E14.5 and E16.5, the cardiac lymphatic vessel network was indistinguishable from controls (Fig. 1C, a, e, f). These results indicate that the critical time requirement for *Tbx1* in cardiac lymphangiogenesis is before E14.5. Thus, *Tbx1* may be required for the migration of lymphatic ECs into the heart or for their coalescence to form cardiac lymphatic vessels.

### Tbx1 and Vegfr3 interact genetically in cardiac lymphatic development

We next tested whether *Tbx1* and *Vegfr3* interact genetically to regulate cardiac lymphangiogenesis. For this, we crossed *Tbx1*^*Cre*/+^ with *Vegfr3*^*flox*/+^ mice (14) and analyzed cardiac lymphatic vessels in E18.5 *Tbx1*^*cre*/+^;*Vegfr3*^*flox*/+^ embryos by whole-mount immunostaining with anti-Lyve-1 antibody. We also generated *Vegfr3*^+/−^ (germline heterozygous) mice by crossing *Vegfr3*^*flox*/+^ mice with ubiquitous Cre-expressing mice. We first analyzed the cardiac lymphatic vessels in *Tbx1*^*Cre*/+^ and *Vegfr3*^+/−^ embryos compared to wild type littermates. We quantified the number, length and width (ratio length/area) of lymphatic vessels in at least five E18.5 embryos per genotype, using the ImageJ software. In *Tbx1*^*Cre*/+^ embryos, we found no differences in the number, length or width of cardiac lymphatic vessels on either the ventral or dorsal surfaces of the heart (Fig. 2A, b, b’ and Fig. 2C, a, a’, b, b’, c, c’) compared to WT controls (Fig. 2A, a, a’). In *Vegfr3*^+/−^ embryos, the number and length of cardiac lymphatic vessels was not different to WT littermates, but both ventral and dorsal lymphatic vessels were moderately dilated (Fig. 2B, b, b’, Fig. 2D, c, c’). We next analyzed compound heterozygous embryos (*Tbx1*^*cre*/+^; *Vegfr3*^*flox*/+^), and found that the cardiac lymphatic network was strongly reduced (Fig. 2A, c) compared to controls (*Tbx1*^*cre*/+^, *Vegfr3*^+/−^, WT), especially on the ventral surface of the heart where, in most embryos, only a few lymphatic vessels were present around the base of the pulmonary trunk and subpulmonary myocardium (arrow in Fig. 2A, c). The dorsal lymphatic network presented as a reduced network of short, dilated vessels that failed to extend to the apex of the heart (arrow in Fig. 2A, c’). Quantitative analysis confirmed a reduction in the number (Fig. 2C, a, a’), length (Fig. 2C, b, b’) and width (Fig. 2C, c, c’) of ventral and dorsal cardiac lymphatic vessels in these mutants. Together, these results demonstrate the existence of a strong genetic interaction between Tbx1 and Vegfr3 in cardiac lymphangiogenesis that affects the number and morphology of the lymphatic vessels.

**Figure 2.**
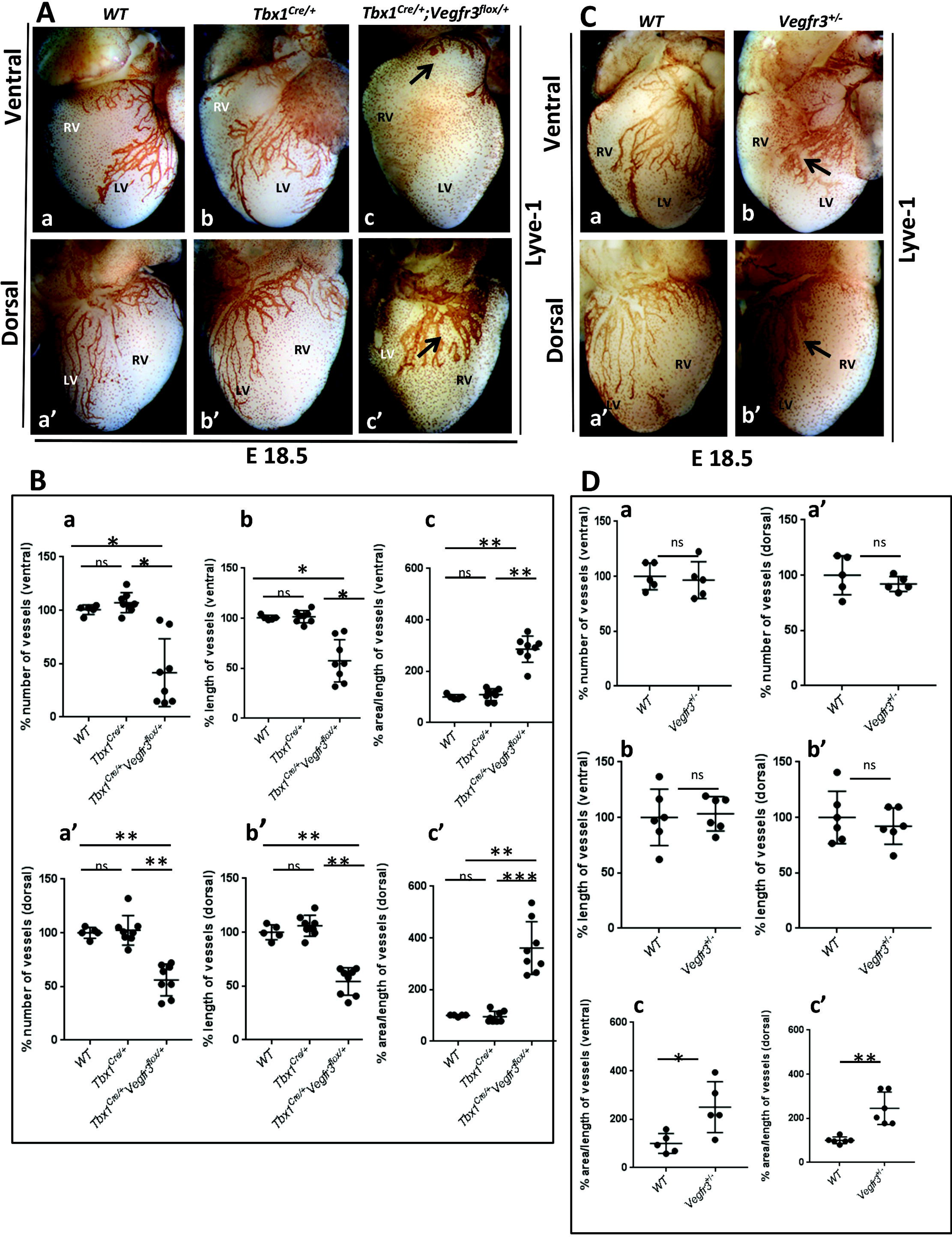
*Genetic interaction Tbx1-Vegfr3 in cardiac lymphatic development*. (A, C) Lyve-1 immunostaining identifies sub-epicardial lymphatic vessels in rappresentative hearts of E18.5 embryos of the genotypes shown (punctate signal derives from Lyve1+ macrophages). (A) Arrows in c, c’ indicate abnormal cardiac lymphatic vessels in compound heterozygous embryos. B) Quantitative analysis revealed a reduced number (a-a’) reduced length (b-b’) and increased width (c-c’) of ventral and dorsal cardiac lymphatic vessels in E18.5 *Tbx1*^*cre*/+^;*Vegfr3*^*flox*/+^ embryos compared to *WT* (a-a’) and *Tbx1*^*Cre*/+^ (control) embryos (b-b’). C) *Vegfr3 haploinsufficiency in cardiac lymphatic development*. Lyve-1 immunostaining reveals mild hypertrophy of cardiac lymphatic vessels in E18.5 *Vegfr3*^*f*+/-^ embryos (b-b’). D) Quantitative analysis (D) shows that number (a-a’) and length (b-b’) of cardiac lymphatic vessels were normal, but they were moderately dilated (c-c’). The cardiac Abbreviations: LV, left ventricle, RV, right ventricle. ∗P<0.05, ∗∗P ≤ 0.01, ∗∗∗P ≤ 0.001. Error bars (SEM).

We also examined the intestinal lymphatic vessels of the same embryos by Lyve-1 immunostaining. Results revealed that also in this tissue, there was lymphatic vessel hypoplasia in *Tbx1*^*cre*/+^; *Vegfr3*^*flox*/+^ embryos, in both the membranous mesentery and in the intestinal wall, compared to controls (*Tbx1*^*cre*/+^, *Vegfr3*^+/−^, WT), although the phenotype was more variable than in the heart (Supplementary Fig. S2). Thus, we conclude that the genetic interaction between *Tbx1* and *Vegfr3* is not limited to cardiac lymphatic development.

### Generation of a *Cre*-inducible *Vegfr3* transgenic mouse

We have hypothesized that lymphatic vessel hypoplasia in *Tbx1* mutants is secondary to reduced Vegfr3 in *Tbx1*-depleted LECs (1) and here we provide evidence of a genetic interaction between the two genes. We next tested whether increasing *Vegfr3* expression in *Tbx1*-expressing tissues, that includes lymphatic ECs, is sufficient to restore normal lymphatic development in single and compound *Tbx1* mutants. We generated a transgenic mouse (named TgVegfr3) that expresses *Vegfr3* upon Cre-induced recombination. For this, we generated a *Vegfr3*-expressing transgene containing a full length murine *Vegfr3* cDNA (Fig. 3A) that we electroporated into E14Tg2A.4 mouse embryonic stem cells (Bay Genomics, CA). We obtained and sequence-verified six clones that had a single integration site and a single copy of the transgene (Fig. 3B, C). Two of these clones were injected into mouse blastocysts. We obtained germline transmission of the transgene in four male offspring. We then characterized the transgenic Vegfr3 protein in cells and in mice. We first tested whether Cre activation of the transgene led to the production of a full length Vegfr3 protein. For this, we co-transfected C2C12 cells, which do not express *Vegfr3*, with the transgene, and a Cre-expressing vector. Western blotting with anti-Vegfr3 antibody performed 24 h after transfection revealed the presence of a protein of the expected length of 195 Kda (Fig. 3D). Furthermore, anti-Vegfr3 immunocytofluorescence performed on the same cells indictated that the transgenic protein localized to the cell membrane (Fig. 3E). In order to confirm that the transgene localized correctly and functioned *in vivo*, we crossed TgVegfr3 mice with *Tbx1*^*Cre*/+^ mice and analyzed expression of Vegfr3 (Fig. 3F) and phosphorylated ERK (pERK, Fig. 3G) in embryos at E15.5. We observed ectopic expression of the transgenic Vegfr3 protein in the tongue (Fig. 3F, a’), heart (Fig. 3F, b’, c’) and mandibular (masseter) muscle (Fig. 3G, a’) in E15.5 *Tbx1*^*cre*/+^; TgVegfr3 embryos but not in *Tbx1*^*Cre*/+^ (control) embryos (Fig. 3F, a-c, Fig. 3G, a). Similarly, pERK was expressed in the masseter muscle of *Tbx1*^*cre*/+^; TgVegfr3 embryos (Fig. 3G, b’) but not in *Tbx1*^*Cre*/+^ (control) embryos (Fig. 3G, b). These results indicate that *Tbx1*^*Cre*^ activates expression of transgenic *Vegfr3* in the endogenous *Tbx1* expression domains and that the transgenic protein is able to activate the MAPK/ERK pathway, a known VegfC/Vegfr3 downstream signalling pathway that promotes cell proliferation.

**Figure 3.**
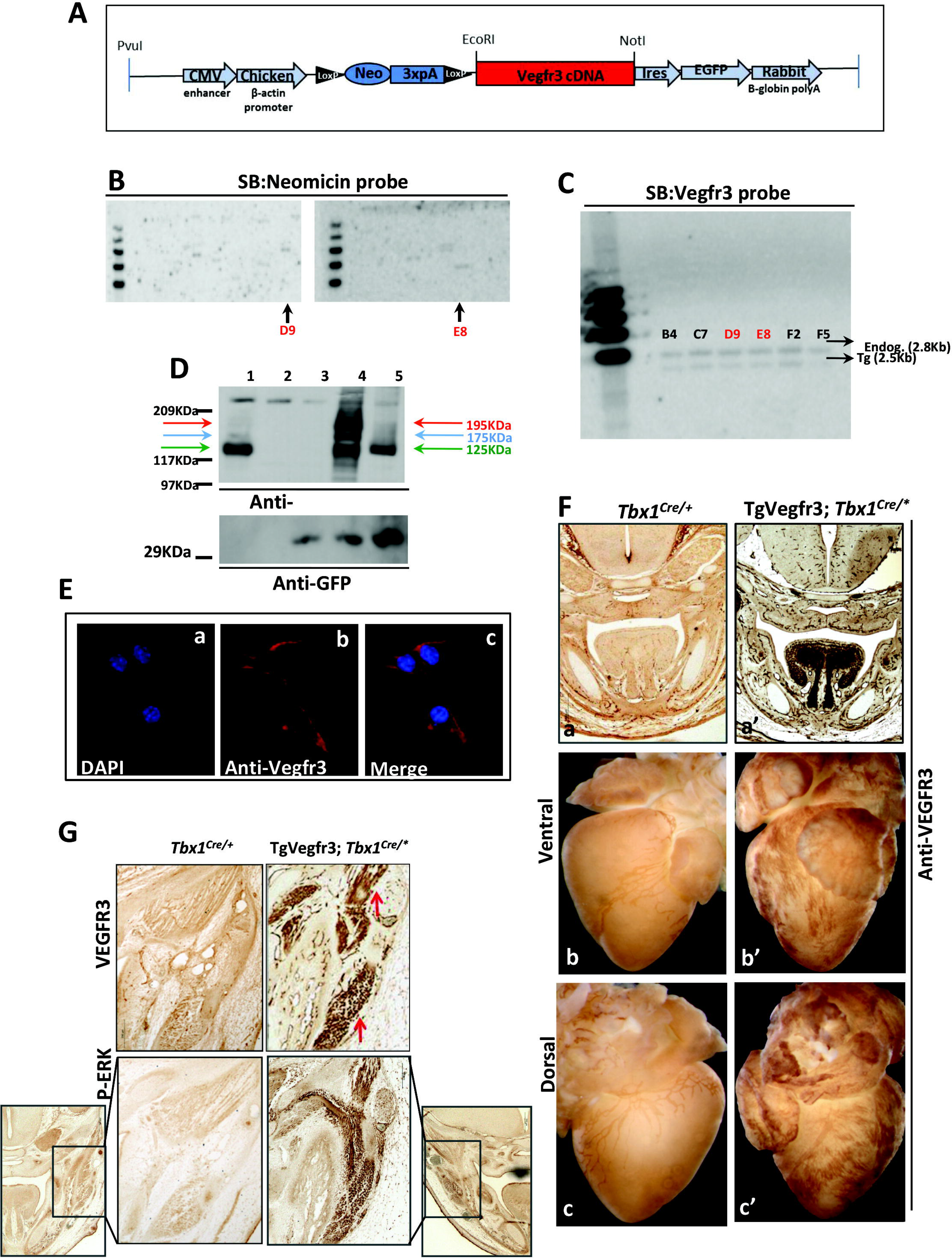
*Generation and in vivo expression of the Cre-inducible Vegfr3 transgene.* (A) Construct used to generate TgVegfr3 transgenic mouse. (B) Southern blotting with the neomycin probe showing 2 out of 22 clones that carried a single copy of the transgene. (C) Southern blotting with the Vegfr3 probe confirmed the identity of the transgene (2.5Kb), compared to the endogenous gene (2.8Kb). Clones D9 and E8 were injected into mouse blastocysts. (D, upper panel) Western blot analysis with anti-VEGFR3 revealed that the transgenic VEFGR3 protein was full length. Proteins were resolved on an 8% SDS-polyacrylamide gel. Lane 1: P19 cells (positive control) express Vegfr3. Lane 2: C2C12 cells (negative control) do not express Vegfr3). Lane 3: C2C12 + empty vector. Lane 4: C2C12 + pCIG2-Vegfr3 plasmid (clone 1). Lane 5: C2C12 + pCIG2-Vegfr3 plasmid (clone 2). In lanes 1 and 4 show three bands corresponding to the full length protein (195KDa, red arrows), the pre-peptide (175KDa, blue arrows) and the proteolytically cleaved form (125 KDa band green arrows). (D, lower panel). The same protein samples probed with anti-GFP. Lanes 3, 4 and 5 show the GFP protein product confirming Cre activated of Vegfr3 and GFP expression. (E) shows Vegfr3 protein localization in C2C12 cells co-transfected with a selected Vegfr3 construct and a construct expressing Cre recombinase. Immunofluorescence was performed 24 hours after transfection: a) DAPI staining of cell nuclei, b) Anti-Vegfr3 staining shows that the transgenic Vegfr3 protein localized correctly to the membrane, c) Merged image of a) and b). (F) Vegfr3 protein production and localization in transgenic mouse. Immunohistochemistry with anti-Vegfr3 antibody on coronal sections of the head of E15.5 *Tbx1*^*Cre*/+^ (a) and TgVegfr3;*Tbx1*^*Cre*/+^ (b) embryos showed ectopic Vegfr3 protein expression in the tongue of TgVegfr3;*Tbx1*^*Cre*/+^ embryos (b). Hearts of E18.5 *Tbx1*^*Cre*/+^ (b, c) and TgVegfr3;*Tbx1*^*Cre*/+^ (b’, c’) embryos showed ectopic Vegfr3 protein expression in the ventral and dorsal myocardium. (G) Immunohistochemistry with anti-Vegfr3 antibody (a, a’) and anti-pERK antibody (b, b’) on coronal sections of the head of E18.5 *Tbx1*^*Cre*/+^ (a, b) and TgVegfr3;*Tbx1*^*Cre*/+^ (a’, b’) embryos showed ectopic expression of both proteins in the masseter muscle in TgVegfr3;*Tbx1*^*Cre*/+^ embryos.

### *Vegfr3* over-expression partially rescues lymphatic defects in *Tbx1-Vegfr3* compound mutants

We tested whether the TgVegfr3 transgene was able to rescue the cardiac lymphatic vessel anomalies observed in *Tbx1*^*cre*/+^; *Vegfr3*^*flox*/+^ embryos. For this, we crossed *Tbx1*^*Cre*/+^ mice with TgVegfr3;*Vegfr3*^*flox*/+^ mice and analyzed cardiac lymphatic vessels in E18.5 embryos with the following genotypes: *Tbx1*^*cre*/+^, *Tbx1*^*Cre*/+^*;Vegfr3*^*flox*/+^, TgVegfr3;*Tbx1*^*cre*/+^;*Vegfr3*^*flox*/+^ (Fig. 4A and Table 1).

**Table 1.** Summary of cardiac lymphatic phenotypes in E18.5 embryos for all genotypes analyzed.*, ratio length/area; #, vessels absent in most embryos.

|  |  | Number of vessels |  | Length of vessels |  | Width of vessels* |  |
| --- | --- | --- | --- | --- | --- | --- | --- |
|  | Number of embryos | ventral | dorsal | ventral | dorsal | Ventral | dorsal |
| WT | 5 | normal | normal | normal | normal | normal | normal |
| *Tbx1*<sup>Cre/+</sup> | 8 | normal | normal | normal | normal | normal | normal |
| *Tbx1*<sup>Cre/+</sup>; *Vegfr3*<sup>fllox/+</sup> | 8 | decreased | decreased | decreased | decreased | Increased | increased |
| TgVegfr3;*Tbx1*<sup>Cre/+</sup>; *Vegfr3*<sup>fllox/+</sup> | 6 | # | decreased | # | normal | # | normal |
| TgVegfr3;*Tbx1*<sup>Cre/+</sup> | 5 | # | normal | # | normal | # | normal |
| *Tbx1*<sup>Cre/lacZ</sup> | 3 | # | # | # | # | # | # |
| TgVegfr3;*Tbx1*<sup>Cre/lacZ</sup> | 4 | # | # | # | # | # | # |
| WT | 5 | normal | normal | normal | normal | normal | normal |
| *Vegfr3*<sup>+/-</sup> | 5 | normal | normal | normal | normal | Increased | increased |

**Figure 4.**
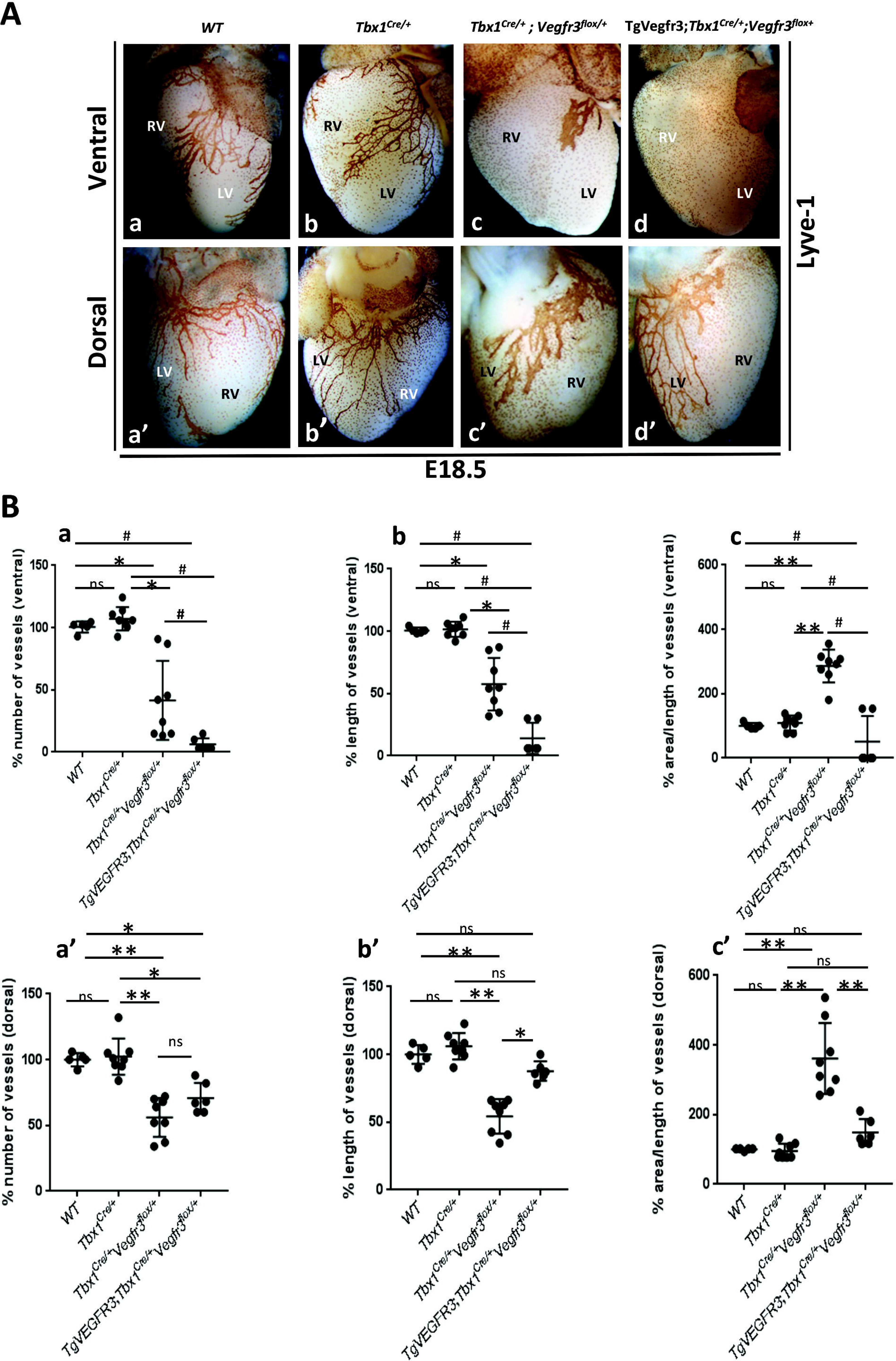
*Vegfr3 expression from the TgVegfr3 transgene partially rescues cardiac lymphatic vessel anomalies in Tbx1-Vegfr3 compound heterozygotes*. (A) Lyve-1 immunostaining identifies cardiac lymphatic vessels in all genotypes shown. Hearts of E18.5 *TgVegfr3*;*Tbx1*^*cre*/+^;*Vegfr3*^*flox*/+^ embryos (d-d’) reveal partial rescue of lymphatic anomalies observed in *Tbx1*^*cre*/+^;*Vegfr3*^*flox*/+^ embryos (c-c’). Quantitative analysis (B) showed rescue of vessel length (b’) and width (c’) but not vessel number (a’) on the dorsal surface of the heart of *TgVegfr3*;*Tbx1*^*cre*/+^;*Vegfr3*^*flox*/+^ embryos compared to *Tbx1*^*cre*/+^;*Vegfr3*^*flox*^ embryos (c, c’). There was no rescue of ventral lymphatic vessels. Abbreviations: LV, left ventricle, RV, right ventricle. ∗P<0.05, ∗∗P ≤ 0.01, ∗∗∗P ≤ 0.001. Error bars (SEM), #, vessels absent in most *TgVegfr3*;*Tbx1*^*cre*/+^;*Vegfr3*^*flox*/+^ embryos (statistical analysis not performed).

Results showed that the transgene was able to rescue partially the cardiac lymphatic phenotype of compound heterozygotes, described in detail above and shown in Fig. 2C, c-c’ and Fig. 4A, c, c’. Specifically, on the dorsal surface of the heart of TgVegfr3; *Tbx1*^*cre*/+^;*Vegfr3*^*flox*/+^ embryos (Fig. 4A, d’), the lymphatic vessel network was similar in appearance to that of *Tbx1*^*Cre*/+^ (control) embryos (Fig. 4A, b’), being composed of mostly thin, well organized vessels, some of which extended to the apex of the heart. Quantitative analysis confirmed that the length and width of the dorsal lymphatic vessels was fully rescued in TgVegfr3; *Tbx1*^*cre*/+^;*Vegfr3*^*flox*/+^ embryos (Fig. 4B, b’, c’). In contrast, the number of lymphatic vessels was not rescued by the transgene (Fig. 4B, a’) suggesting that their growth is regulated by *Tbx1*, consistent with our previously published findings (1). On the ventral surface of the same hearts, there was no appreciable rescue, in fact three out of five embryos lacked ventral lymphatic vessels, precluding a statistical analysis. These findings suggested that lymphatic vessel hypertrophy in *Tbx1*^*cre*/+^; *Vegfr3*^*flox*/+^ embryos was due to reduced dosage of *Vegfr3* in ECs, and that the transgene was able to compensate for this in *TgVegfr3; Tbx1*^*Cre*/+^*;Vegfr3*^*flox*/+^ embryos. Furthermore, as the number of lymphatic vessels was not rescued by the transgene, The different effects of the transgene on the dorsal versus ventral surface of the heart was puzzling initially, but became more clear when we analyzed TgVegfr3; *Tbx1*^*Cre*/+^ embryos. In these mutants, the dorsal cardiac lymphatic vessels were indistinguishable from WT and *Tbx1*^*Cre*/+^ embryos while the ventral surface of the heart lacked lymphatic vessels (Supplementary Fig. S3). This suppressive effect of the transgene on ventral cardiac lymphangiogensis likely accounts for it’s inability to rescue the ventral lymphatic hypoplasia caused by combined Tbx1-Vegfr3 haploinsufficiency..

As the transgene was quite effective at rescuing lymphatic anomalies on the dorsal surface of the heart, we tested whether it modified the cardiac lymphatic phenotype of *Tbx1* homozygous mutants (TgVegfr3;*Tbx1*^*Cre/lacZ*^), with negative results (Table 1 and Supplementary Fig. S3).

In conclusion, we propose a model where Tbx1 and Vegfr3 interact in lymphatic ECs to establish a reciprocal gene dosage equilibrium that determines the correct number and morphology of sub-epicardial lymphatic vessels. The absence of lymphatic vessels on the ventral surface of the heart of different genetic mutants used in this study suggests that the Tbx1-Vegfr3 interaction is required prior to, or at the onset of cardiac lymphangiogenesis.

## DISCUSSION

Our previous study indicated that *Tbx1* regulates lymphatic development through the VegfC-Vegfr3 axis. In this study, we have demonstrated a strong genetic interaction between *Tbx1* and *Vegfr3* that is critical for cardiac lymphangiogenesis and involves two distinct phenotypes, namely, the number of lymphatic vessels and lymphatic vessel morphology. *In vivo* rescue experiments indicated that these different aspects of the cardiac lymphatic phenotype are regulated independently by the two genes, with Tbx1 regulating the number of lymphatic vessels and Vegfr3 their morphology. We also found that for cardiac lymphatic development *Vegfr3* is haploinsufficient, a feature that, to our knowledge, has not been reported before.

Rebalancing *Vegfr3* gene dosage in *Tbx1*-deficient cells was not sufficient for normal cardiac lymphatic development, suggesting that there may be other effectors of Tbx1 function in lymphatic ECs. However, Tbx1 may also have a developmental function in EC progenitors that cannot be compensated by enhanced *Vegfr3* expression. This hypothesis arises from the observation, reported here, that *Tbx1*-expressing cells and their descendants contribute extensively to the cardiac lymphatic network, and they are present in the earliest cardiac lymphatic vessels. Recently, it has been demonstrated that cardiac lymphatic vessels derive from venous and non-venous EC progenitors (2). In the future, it would be interesting to determine whether there is a contribution from the secondary heart field, where *Tbx1* is highly expressed, and where *Tbx1*-expressing multi-potent progenitors give rise to ECs that populate the aortic arch arteries and cardiac outflow tract (10), as well as to cardiomyocytes and branchiomeric muscles (15)(16).

We were intrigued by the different response of the ventral and dorsal surfaces of the heart to altered *Vegfr3* gene dosage. We have shown here that the genetic interaction between *Tbx1* and *Vegfr3* is especially evident on the ventral surface of the heart (Fig. 2A-c). Furthermore, the capacity of TgVegfr3 to rescue lymphatic anomalies in *Tbx1-Vegfr3* compound heterozygotes was limited to the dorsal surface of the heart (Fig. 4d’). Finally, over-expression of *Vegfr3* from the TgVegfr3 transgene had a strong effect on the ventral but not dorsal cardiac lymphatic vessels (Fig. 4e, 4e’). Note that in all three cases, a single copy of *Tbx1* (*Tbx1*^*cre*/+^) was inactivated, which alone had no effect on cardiac lymphatics. Together, these results suggest that *Tbx1* mutation sensitizes the ventral surface of the heart to altered *Vegfr3* gene dosage (mild lymphatic hypertrophy in *Vegfr3*^+/−^ mutants affects ventral and dorsal cardiac lymphatic vessels (Fig. 2C).

What might be the basis of this differential sensitivity? It is not likely to be linked to differences in activation of the transgene as this appeared to be uniform in the dorsal and ventral heart (Fig. 3F, b-c). In fact, the differential response was also evident in the absence of the transgene, in *Tbx1*^*cre*/+^;*Vegfr3*^*flox*/+^ compound heterozygotes. Other possibilities include, i) differences between lymphatic EC progenitors; *Tbx1* may be more highly expressed in ventral progenitors, ii) differences in vessel growth; *Tbx1* may be more highly expressed in ventral cardiac lymphatic vessels, and when it is absent they fail to grow, iii) differences in the local environment. Considering the first possibility, studies of early cardiac lymphatic vessel development have shown that they derive from an extra-cardiac source of Prox1+/Lyve1+ lymphatic ECs that are of mainly venous origin (cardinal vein), which reaches the heart at E12.5-E13 from the mediastinum on the dorsal surface of the heart and from the cardiac outflow tract on the ventral surface (17)(5)(2) (18). Whether these invading lymphatic ECs derive from different progenitors has not been reported. Here, we have shown that *Tbx1*-expressing cells and their descendants contribute extensively to both ventral and dorsal lymphatic networks suggesting that differential *Tbx1* expression in EC progenitors, or their descendants, does not account for the differential ventral ≫ dorsal sensitivity. Considering the second possibility, from E14.5, lymphatic vessels on the ventral and dorsal surfaces of the heart grow in a base-to-apex direction, extending more rapidly over the dorsal surface (2). If *Tbx1* were more highly expressed in ventral lymphatic vessels, or if there were a critical *Tbx1* function or Tbx1-Vegfr3 interaction in ventral lymphatic vessels, this might preferentially stall the growth of the ventral lymphatic network. Our results show that *Tbx1* is expressed in both ventral and dorsal lymphatic vessels at E14.5 (Fig. 1B-b’, 1B-d’), but this does not exclude functional differences between the dorsal and ventral networks. Considering the third possibility, several studies have shown that the local molecular environment can profoundly influence Vegfr3 function in the context of blood vessel development. For example, it has been demonstrated that *Vegfr3* may promote or antagonize angiogenesis through mechanisms that are Notch-dependent and VEGF-ligand-independent (19)(20). These mechanisms may also depend upon *Tbx1* dosage. As yet, there are no studies that address the issue in lymphatic vessels, but it is certainly possible that local differences in the molecular milieu on the ventral versus dorsal surfaces of the heart determine a greater sensitivity to the ventral network to altered Vegfr3 dosage. Finally, increased *Vegfr3* expression in the endogenous domain or ectopic *Vegfr3* expression may negatively affect cardiac lymphangiogenesis. We attempted to address this by generating *TgVegfr3;Tie2*^*Cre*/+^ embryos, but we did not recover any embryos with the desired genotype at E14.5, indicating that forcing expression of *Vegfr*3 in all endothelial cells causes lethality prior to the onset of cardiac lymphangiogenesis. Evidently, reducing endogenous *Vegfr3* expression in *TgVegfr3;Tbx1*^*Cre*/+^*;Vegfr3*^*flox*/+^ embryos is not sufficient to rescue cardiac lymphatic hypoplasia caused by expression of the transgene. This suggests that ectopic *Vegfr3* expression driven by *Tbx1*^*Cre*^, for example in cardiomyocytes of the right ventricle, may be responsible for lymphatic hypoplasia in these mutants, perhaps by sequestering the ligand and thus reducing its availability to lymphatic ECs.

Intriguingly, recent clinical studies have linked *VEGFR3* mutations to congenital heart disease (CHD), and the 22q11.2 deletion to lymphatic problems, encouraging speculation about a genetic interaction between *TBX1* and *VEGFR3* in humans. In the first study, Jin et al. showed for the first time that *VEGFR3* might play a critical role in heart development (21). In this study, exome sequencing of 2871 probands with CHD revealed dominant *VEGFR3* mutations in 2.3% of cases of Tetralogy of Fallot (TOF), the most common CHD associated with 22q11.2DS, for which *TBX1* is the major disease gene. In the second study, Unolt et al. reported the presence of primary lymphedema in 4/1600 (0.025%) patients with 22q11.2DS (22). Prior to this, only a single case report had documented lymphatic abnormalities in a patient diagnosed with DiGeorge syndrome (23). The finding of primary lymphedema in 22q11.2DS patients supports the hypothesis of a TBX1-VEGFR3 interaction, through which *TBX1* heterozygosity predisposes to lymphatic anomalies by disrupting the VEGF/C-VEGFR3 signalling pathway. Considering the recent clinical findings and supporting data from mouse models of 22q11.2DS, it would be interesting to determine the frequency of deleterious *VEGFR3* variants in 22q11.2DS patients, and establish whether this leads to more severe types of CHD.

## MATERIAL AND METHODS

### Mouse lines and tissues

The following mouse lines were used: *Tbx1*^*lacZ*/+^ (8), *Tbx1*^*flox*/+^ (24), *Tbx1*^*Cre*/+^ (25), Tie2Cre (26), *Vegfr3*^*flox*/+^ (14), R26R (12), *Rosa*^*mTmG*^ (13), TgCAGG-CreER™ (27). Genotyping of mice was performed according to the original reports. Activation of CAGG-CreER™ was performed with a single intraperitoneal injection of Tamoxifen (Sigma) at 75 mg/kg body weight at one of various time points between gestation days 10.5 and 16.5.

### Salmon-gal staining, immunohistochemistry and immunofluorescence

β-Gal activity was revealed by salmon-gal staining. Embryonic hearts at E14.5, E15.5 and E18.5 were isolated and fixed for 30 min in glutaraldehyde (0.2%), paraformaldehyde (2%), EGTA (5 mM), magnesium chloride (2 mM) and phosphate buffer a pH 7.3 (0.1M), then washed three times (20 mins per wash) in a solution containing sodium deoxycholate (0.1%), NP40 (0.2%), magnesium chloride (2 mM) and phosphate buffer pH 7.3 (0.1M). Hearts were stained in a solution containing salmon-gal (1mg/mL) and NBT (0.4 mM) in washing solution in the dark at 37°C with gentle rotation until the desired staining was achieved. Immunostaining was performed using the following primary antibodies: rat anti–Vegfr3 (eBioscience), rabbit anti-Lyve1 (Abcam), rabbit anti-GFP (Invitrogen), mouse anti-pERK (Cell Signalling), diluted 1: 400, and secondary antibodies: HRP-conjugated goat anti-rat IgG (Kirkegaard & Perry Laboratories), HRP-conjugated anti-rabbit IgG (GE Healthcare), diluted 1:200. Fluorescent antibodies were visualized using Vectashield imaging medium (Vector Laboratories). Non-fluorescent antibodies were visualized using DAB (Vector Laboratories). Whole-mount specimens were photographed using a dissecting microscope (Stemi 2000; Carl Zeiss, Inc.) equipped with a camera (Axiocam; Carl Zeiss, Inc.) and the manufacturer’s acquisition software. Images were acquired at 28x magnification.

### Quantitative analysis of lymphatic vessel anomalies

We analyzed a minimum of 5 and a maximum of 8 embryos per genotype in all experiments. Three-dimensional images were digitally reconstructed from *z* stacks. For each image, we manually counted all the lymphatic vessels on the dorsal and ventral surfaces of the heart, and we measured the length and area of each vessel. We then calculated the ratio length/area to determine the width of each single vessel. Quantitative analysis was performed using the ImageJ software, (www.uhnresearch.ca/facilities/wcif/imagej).

### Statistical analysis

We used the non-parametric Kruskall-Wallis test for the analysis of lymphatic vessels in all experiments with a single exception, namely *Vegfr3*^+/−^ vs WT embryos, for which the Mann-Whitney U test was used. We first calculated the mean number of vascular features (number, length, area/length vessels) for each embryo. We then calculated the mean value for the group (same genotype). The latter value was used for the statistical analysis.

### Generation of transgenic mouse embryonic stem cells

The starting vector was the pCIG2 plasmid (28) without the nuclear localization sequence of EGFP-CDS that consists of a CMV-IE enhancer, chicken β-actin promoter, MCS, IRES-GFP and rabbit β-globin polyA. A loxP-flanked neomycin resistance cassette was cloned downstream of the β-actin promoter followed by a full-length mouse *Vegfr3* cDNA (GeneCoepia). The linearized Vegfr3 transgenic construct, named TgVegfr3, was electroporated into feeder-free E14Tg2A.4 mouse ES cells using the ECM 630 Electro Cell Manipulator System (BTX). Twentyfour hours post-electroporation, the cells were selected in G418 at a final concentration of 250μg/ml. ES cell clones were picked 12 day after plating.

### Identification of correctly targeted ES cell clones by southern blotting

Two probes were generated for southern blotting experiments. The first probe distinguished the transgene from the endogenous *Vegfr3* gene. It was generated by PCR amplifying a segment of exon 13 of the *Vegfr3* cDNA, using the following primers: pF 5’CTTAGAAGGCCAGTCCGTGC 3’, pR 5’CCTGCACGGACAGGTACTTC 3’. BamHI digestion of genomic DNA from ES clones generated a 2.5Kb fragment in the presence of the transgene and a 2.8Kb fragment in the endogenous *Vegfr3* gene. To identify clones with a single insertion of the transgene, a second probe was generated as follows; a region of the neomycin resistance cassette was amplified by PCR using the primers: pF 5’ GCACAACAGACAATCGGCTG 3’, pR 5’GATACTTTCTCGGCAGGAGC 3’. This probe, which was designed upstream the first BamHI cutting site in the linearized TgVegfr3 construct, localized within the neomycin-resistance cassette. The first BamHI cutting site upstream of the region amplified localized to genomic DNA, and thus its’ position depended upon the insertion site of the electroporated TgVegfr3 construct. Thus, the size of the fragment recognized by this probe varied from clone to clone, and within in the same clone, from insertion to insertion, in the case of multiple integrations of the transgene. ES cell clones containing a single copy of TgVegfr3 were analyzed by PCR to confirm that the entire *Vegfr3* cDNA was present, using the primers: pF 5’CGACGAATTCGGTACCATGCAG 3’, pR 5’GCACGGACTGGCCTTCTAAG 3’. These primers amplified a region extending from 16 bp upstream of the ATG to 20 bp downstream of the BamHI site. A 1741 bp fragment was expected.

### Generation of transgenic mice

Two correctly targeted ES cell clones were selected for microinjection into mouse blastocysts. Four males chimeras were obtained. These males were bred with black C57BL/6 female mice and three germ line transmissions were obtained. These founders were crossed into the C57BL/6 strain. Mice were genotyped by PCR using DNA extracted from tail biopsies, using following primer pairs: F-5’-ATCGACCTGGCAGACTCCAA-3’; R-5’-GAAAACTGCGATGACGCCAGT-3’. PCR products were separated on 2% agarose gels.

### Western blot analysis

Transfected C2C12 cells were collected 24h after transfection, washed with PBS and lysed in lysis buffer (TrisHCl 20mM pH7.4, NaCl 100mM, 10mM MgCl2, NP-40 1X, Glycerol 10%, proteases inibitors). Denatured proteins were separated by SDS-PAGE, transferred to Immuno-Blot PVDF Membrane (BioRad) for protein blotting, Membranes were blocked for 1 h at RT in TBST (150 mM NaCl, 10 mM Tris-HCl, pH 7.4, and 0.05% Tween) and powdered milk. The membranes were incubated overnight at 4°C in primary antibodies diluted in TBST-5% milk, then for 1 h at RT with horseradish peroxidase-conjugated secondary antibodies (diluted in TBST-5% milk). Protein binding was detected by ECS (Amersham) using Hyperfilm (Amersham). The molecular masses of proteins were estimated relative to the electrophoretic mobility of the co-transferred, pre-stained protein marker All Precision Blue (Biorad). The primary antibodies used were; rat anti-Vegfr3 (eBioscience), rabbit anti-GFP (Sigma-Aldrich).

### Immunofluorescence on C2C12 cells

C2C12 were plated into 6-well plates on coverslips. The next day, cells were co-transfected with the TgVegfr3 construct and Cre recombinase vector. Immunocytofluorescence was performed 24 hours after transfection. Cells were washed and fixed with 3% paraformaldehyde/PBS for 20 min at RT, permeabilized with 0.1% Triton X-100/PBS for 25 min and blocked with 1% BSA/PBS for 45 min. Cells were incubated with anti-Vegfr3 antibody (eBioscience) diluted 1:100 in 0,1% BSA/PBS for 3 hours, washed 3 times with PBS, incubated for 1 hr with secondary antibody AlexaFluor anti-rat 494nm (Invitrogen) 1:200 in PBS. The fluorescent antibody was visualized using imaging medium (Vectashield with DAPI, Vector Laboratories). For visualization, the coverslip was positioned on a slide and images were acquired using confocal microscopy.

## ACKNOWLEDGEMENTS

We are grateful for the support provided by the Integrated Microscopy Core and the Animal Facility at the Institute of Genetic and Biophysics ‘ABT’/CNR, Naples. *Vegfr3*^*flox*^ mice were kindly provided by Dr. Kari Alitalo.

## SUPPLEMENTARY FIGURE LEGENDS

Supplementary Figure S1

*Tbx1-expressing cells and their descendants populate cardiac lymphatic vessels*. β-gal staining of E18.5 *Tbx1*^*cre*/+^; R26R hearts (a) and in transverse sections of the heart (a’) shows that *Tbx1* is expressed in cardiac lymphatic vessels, identified by anti-Vegfr3 immunohistochemistry in c, c’ c’’, and in myocardium of the right ventricle.

Supplementary Figure S2

*Genetical interaction Tbx1-Vegfr3 in intestinal lymphatic development*. Lyve-1 immunostaining identifies lymphatic vessels in the intestinal tract of E18.5 embryos of all genotypes shown. White arrows indicate lymphatic vessels in the intestinal wall and in the membranous mesentery.

Supplementary Figure S3

*Forced expression of Vegfr3 suppresses ventral cardiac lymphangiogenesis on a Tbx1 heterozygous background*. Lyve-1 immunostaining identifies cardiac lymphatic vessels in all genotypes shown. A. Hearts of E18.5 *TgVegfr3*;*Tbx1*^*Cre*/+^ embryos show normal cardiac lymphatic vessels on the dorsal surface and absence of these vessels on the ventral surface. B. *Tbx1* homozygous embryos (a, a’) lack cardiac lymphatic vessels. Increasing *Vegfr3* expression in *Tbx1*-deficient cells via *Tbx1*^*Cre*^-mediated activation of the transgene does not modify this phenotype (b, b’).

