## Supplementary figures and images for "*Tbx1* interacts genetically with *Vegfr3* to regulate cardiac lymphangiogenesis in mice"

### Supplementary Figure S1

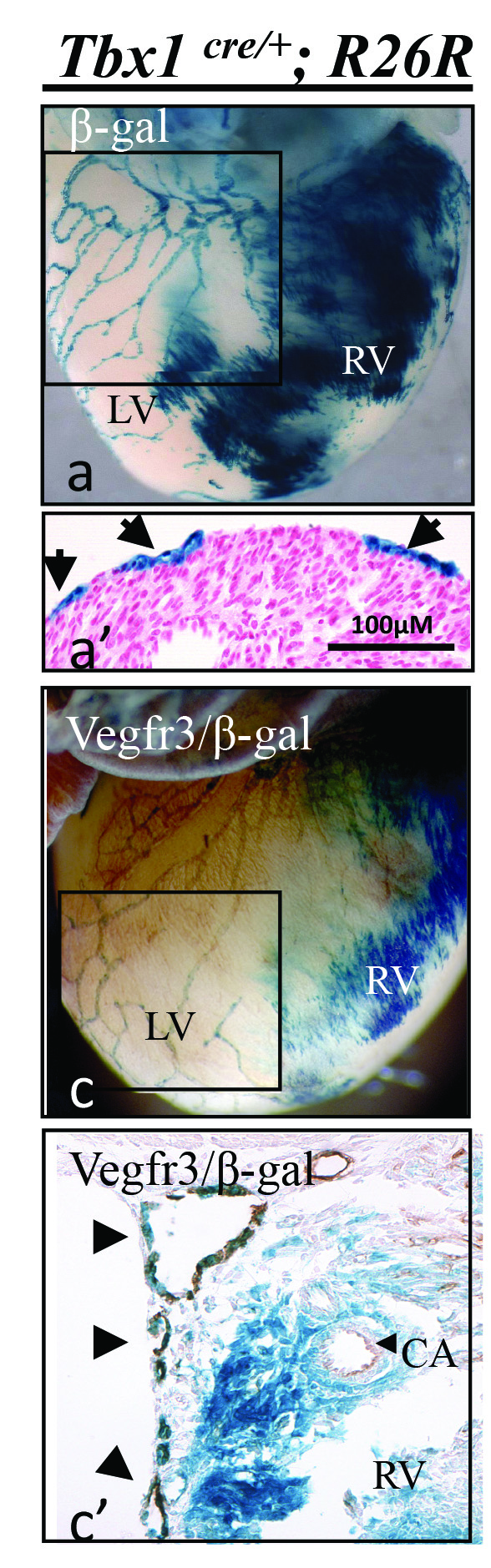

### Supplementary Figure S2

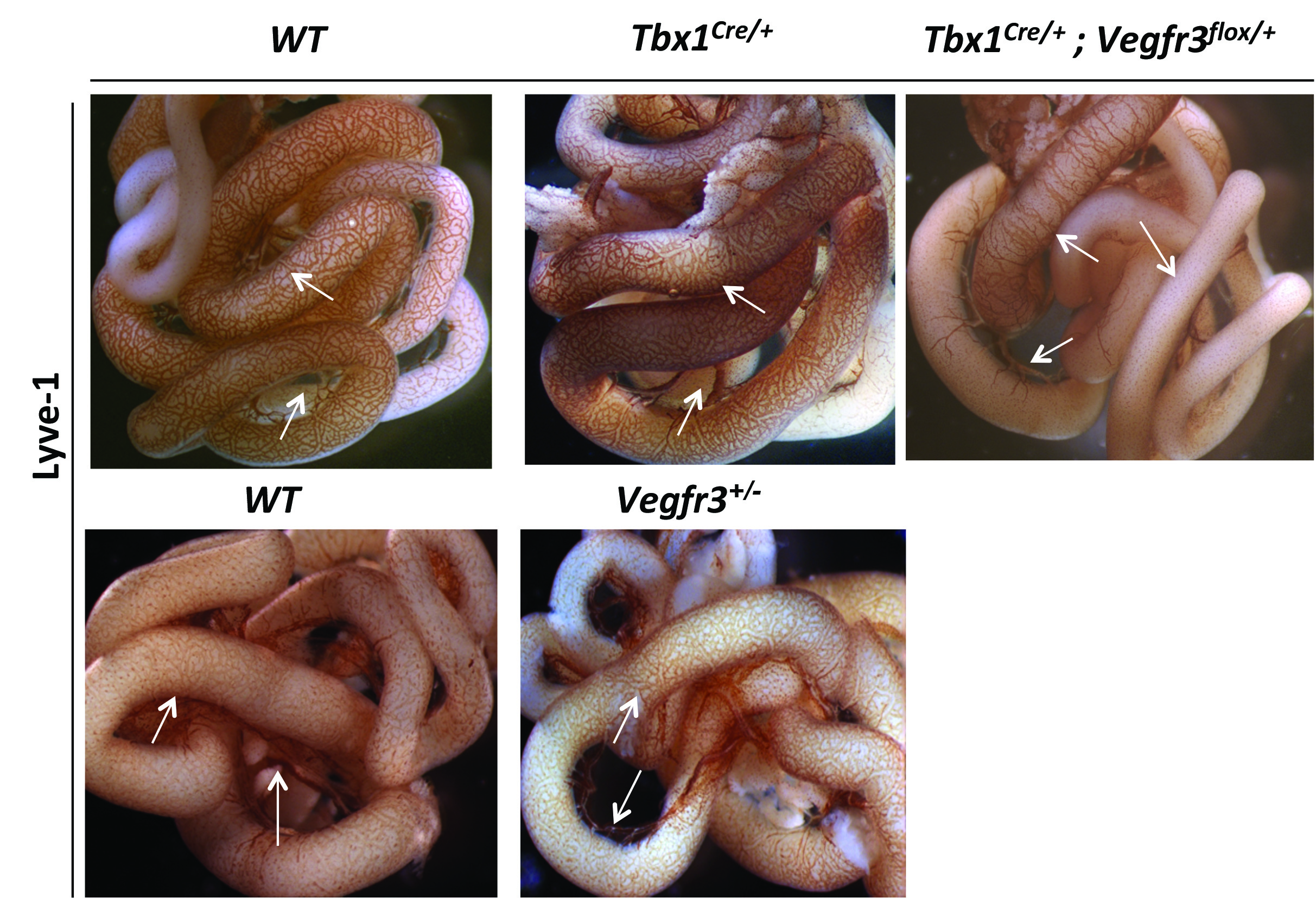

### Supplementary Figure S3

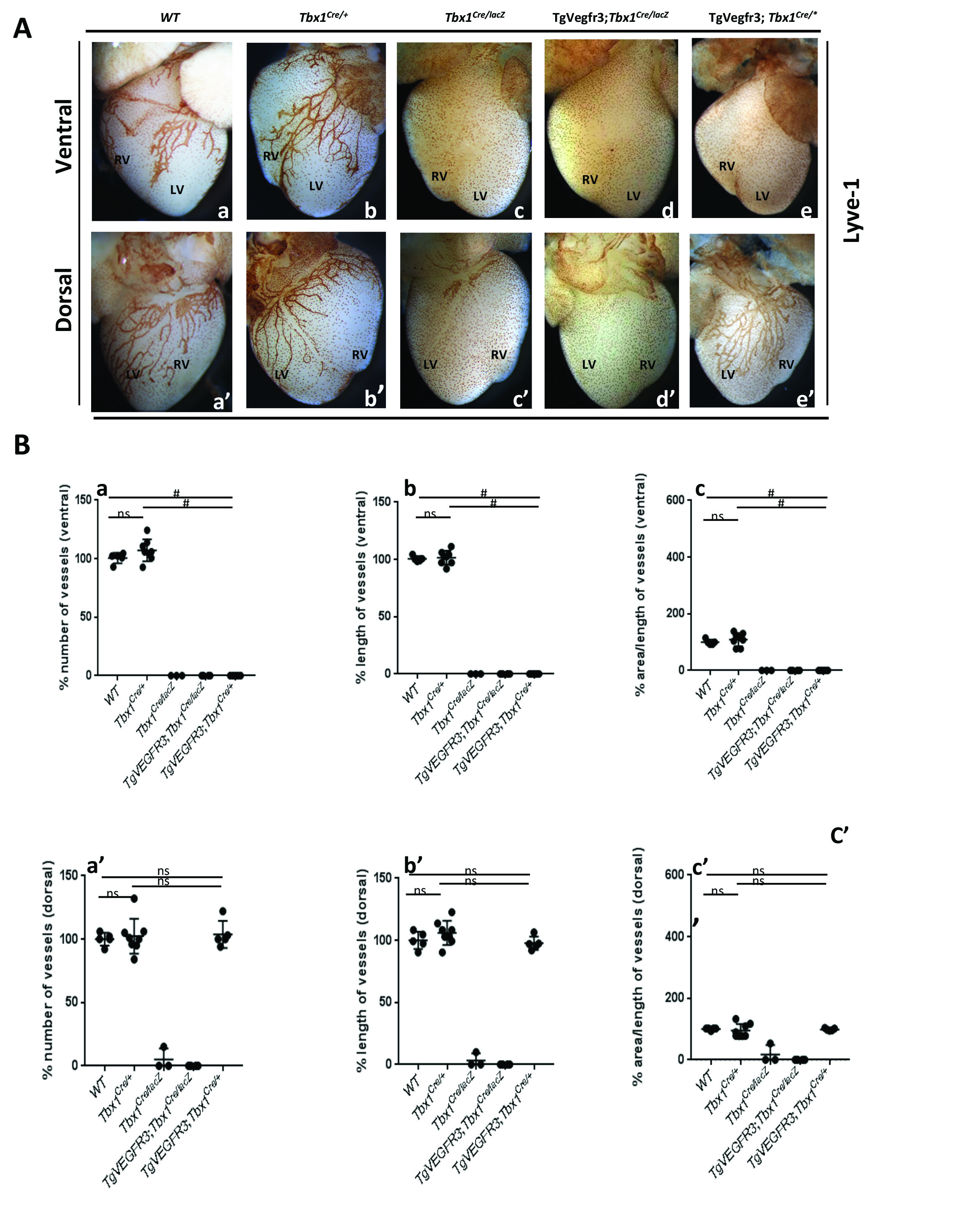
